# Structural correlates of disease-associated CTSC variants in Papillon–Lefèvre syndrome

**DOI:** 10.64898/2026.11.18.767647

**Authors:** Filipe Vono, Andre Naufel, Valeria Santos

## Abstract

**Objective:** To quantify structural features of disease-associated CTSC missense substitutions and evaluate a structural rule for research prioritization in Papillon–Lefèvre syndrome.

**Design:** We compared 16 ClinVar pathogenic or likely pathogenic missense substitutions with 234 recurrent gnomAD substitutions in the mature cathepsin C structure (PDB 1K3B). An expanded exploratory analysis included 35 further literature-indexed substitutions. We audited the documentation used to date the historical holdout and compared AlphaGenome splice scores with a consequence-only baseline.

**Results:** Median predicted destabilization was 2.209 versus 0.763 kcal/mol (AUC 0.764, 95% CI 0.612–0.897). Catalytic proximity was observed in 4/16 versus 9/234. The expanded comparison gave AUC 0.740. Sixteen of 31 substitutions in the historical post-2015 subset had references dated through 2015. For noncoding variants, AlphaGenome splice scores and the consequence-only baseline both obtained AUC 1.000 on 12 positive and 105 negative ClinVar records.

**Conclusions:** Predicted destabilization was associated with ClinVar label status in this dataset. The literature chronology and noncoding consequence composition limit claims of predictive validation.

## 1. Introduction

Papillon-Lefèvre syndrome (PLS) is a rare autosomal recessive disorder characterized by severe early-onset periodontitis leading to premature loss of primary and permanent dentition, accompanied by palmoplantar keratoderma (Toomes et al., 1999; Hart et al., 2000). PLS, Haim-Munk syndrome, and a subset of nonsyndromic aggressive periodontitis cases are caused by biallelic loss-of-function variants in CTSC, which encodes lysosomal cathepsin C (dipeptidyl peptidase I, DPPI; UniProt P53634, 463 amino acids). Mature DPPI functions as a homotetramer assembled from an exclusion domain (partly resolved from residues 25–143 in 1K3B), a heavy chain (231–394) and a light chain (395–463) following cleavage of the signal peptide (1–24) and the intervening propeptide, with a catalytic triad formed by Cys258, His405, and Asn427 (Turk et al., 2001). Because DPPI activates neutrophil serine proteases (neutrophil elastase, cathepsin G, proteinase 3), loss of DPPI activity may alter periodontal host defense (Pham et al., 2004).

Case reports and catalogues have described structural interpretations of CTSC substitutions (Nagy et al., 2014; Moghaddasian et al., 2014; Chitsamankhun et al., 2024). We asked which structural features distinguished disease-associated substitutions from recurrent population substitutions, and how a simple rule behaved under internal cross-validation and on a separate literature-derived subset.

We also examined whether nucleotide-level predictions could complement protein-level scores in selected splice-related variants.

## 2. Materials and Methods

### 2.1 Analysis plan and grouping

The repository records the analysis plan in PRE_REGISTRO.md at commit 574901d, preceding the first public commit of results. The plan specified the distance cutoffs, rule forms and intended temporal split. Later propeptide, Asp25, expanded predictor and AlphaGenome comparisons are exploratory. A subsequent audit traced references for all 35 literature-indexed candidates and assessed grouped resampling.

The unit of analysis was a protein substitution. The primary comparison included only ClinVar pathogenic or likely pathogenic (P/LP) substitutions with 1K3B coverage. Recurrent population comparators had gnomAD allele count ≥ 2 after excluding both ClinVar P/LP and literature-indexed candidates. The comparator pool was held fixed across analyses. Population recurrence does not establish benign status.

The expanded exploratory analysis used the original 35 literature-indexed positives alongside ClinVar P/LP records. These candidates were originally selected by text matching, reference-residue verification, frequency < 10^−4^ and absence of a recorded ClinVar conflict. We audited references and dates using the review table and primary reports where directly accessible. The supplement distinguishes verified primary evidence, indexed findings and unresolved source links. The dated reference is an observed source, which may postdate the earliest report. No change was made to clinical classifications.

### 2.2 Structural descriptors and stability

We used the human cathepsin C biological assembly (PDB 1K3B, 2.15 Å), mapping 352 resolved residues per subunit to UniProt P53634. We computed minimum heavy-atom distances to the catalytic triad (Cys258, His405 and Asn427), nearest other subunit and chloride ions, as well as relative solvent accessibility and annotated disulfide and glycosylation sites. ThermoMPNN estimated ΔΔG from a coordinate input combining the three mature chains of a subunit and from the full-length AlphaFold DB model AF-P53634-F1 (Dieckhaus et al., 2024). Positive scores indicate predicted destabilization. The calculation on the mature subunit estimates substitution effects in the input structure and does not estimate tetramer binding or precursor processing energetics.

The four-term rule was positive if any of four criteria were met: distance to the catalytic triad < 4.0 Å, distance to another subunit < 4.0 Å, substitution at an annotated disulfide residue, or ΔΔG ≥ the selected threshold. The two-term rule retained catalytic distance and ΔΔG. In each held-out fold, the training subset selected a threshold by maximizing Youden’s index on a grid from 0 to 3.00 kcal/mol in 0.25-kcal/mol increments. The modal training-fold threshold was then used for the literature holdout. An exploratory check repeated the cross-validation with all substitutions at the held-out residue removed from training. The figure showing unclassified substitutions uses a fixed threshold of 1.5 kcal/mol.

### 2.3 Predictor comparisons and noncoding analysis

Exploratory AUCs for ESM-1b, REVEL, MetaRNN and CADD were calculated using records with available GRCh38 dbNSFP annotations obtained via MyVariant.info. AlphaMissense and ThermoMPNN AUCs in the main analysis used the larger structure-mapped set. Scores from differently covered subsets were reported descriptively. No paired ranking across all predictors was performed.

AlphaGenome v2 outputs were summarized as maximum absolute scores within K562, HL-60 and skin fibroblast conditions, retaining CTSC-associated and unassigned tracks as implemented in the analysis script. K562 was the primary condition selected using the presence of a CTSC RNA-seq signal relative to ALB. We contrasted ClinVar pathogenic/likely pathogenic with benign/likely benign labels among 152 scored noncoding records. As an annotation-only comparator, we coded entries reported as splice donors or acceptors as 1 and other entries as 0. The same variants were analysed across the three conditions. Scoring of c.757G>A (p.Ala253Thr) was an illustrative retrospective comparison with published functional evidence.

Continuous and categorical comparisons used two-sided Mann–Whitney U and Fisher exact tests, respectively. The original analysis used Wilson intervals for proportions and stratified bootstrap intervals for AUCs, resampling substitutions. The ClinVar-only analysis uses 2,000 stratified bootstrap replicates with a fixed seed. Reported p values are nominal. Co-occurrence of substitutions at the same residue and the small positive propeptide sample constrain inference.

## 3. Results

### 3.1 Structural correlates and exploratory comparisons

Among the 16 ClinVar P/LP and 234 population comparators, median predicted 1K3B destabilization was 2.209 versus 0.763 kcal/mol (AUC 0.764, bootstrap 95% CI 0.612–0.897, p = 0.000412). At 1.5 kcal/mol, 11/16 positives and 62/234 comparators reached the threshold (nominal p = 0.000833). Catalytic proximity occurred in 4/16 versus 9/234 (nominal p = 0.00571). Substitutions at disulfide residues occurred in 1/16 versus 5/234 (nominal p = 0.330).

In the expanded analysis (51 versus 234), the mature-structure ΔΔG AUC was 0.740 (bootstrap 95% CI 0.657–0.813). The expanded comparison showed 8/51 versus 9/234 near the catalytic triad and 5/51 versus 5/234 at disulfide residues. Other structural comparisons, including Asp25 distance, were exploratory. The accompanying structural figures are supplementary and describe the historical expanded analysis.

For the region annotated as propeptide in the consolidated table, the full-length AlphaFold model yielded AUC 0.946 (3 positive substitutions and 62 population comparators). G139R was mapped to 1K3B despite being assigned to this region in the table. Restriction to variants without 1K3B coverage gave AUC 0.922 (Y168C and L196P versus 58 comparators). The estimated AUCs are sensitive to the limited number and definition of positive substitutions. AlphaMissense scored Y168C at 0.230 while its full-length-model ThermoMPNN estimate was 2.352 kcal/mol.

The ClinVar-only AlphaMissense AUC was 0.769 (bootstrap 95% CI 0.647–0.873). In the expanded analysis its AUC was 0.819. On the score-available subsets reported by the original benchmark, ESM-1b, REVEL, MetaRNN and CADD gave AUCs of 0.820, 0.801, 0.790 and 0.783, respectively (reported coverage: 30 positives and 233 recurrent comparators). These are descriptive performance estimates with different coverage from the main analysis.

### 3.2 Threshold selection and literature holdout

The four-term rule gave 15/20 positive calibration substitutions and 174/234 negative population comparators in leave-one-variant-out cross-validation (Figure 4). Thresholds ranged from 2.25 to 2.75 kcal/mol; the mode was 2.25. Leaving out all substitutions at each held-out residue gave 15/20 and 175/234, respectively.

**Figure 1.**
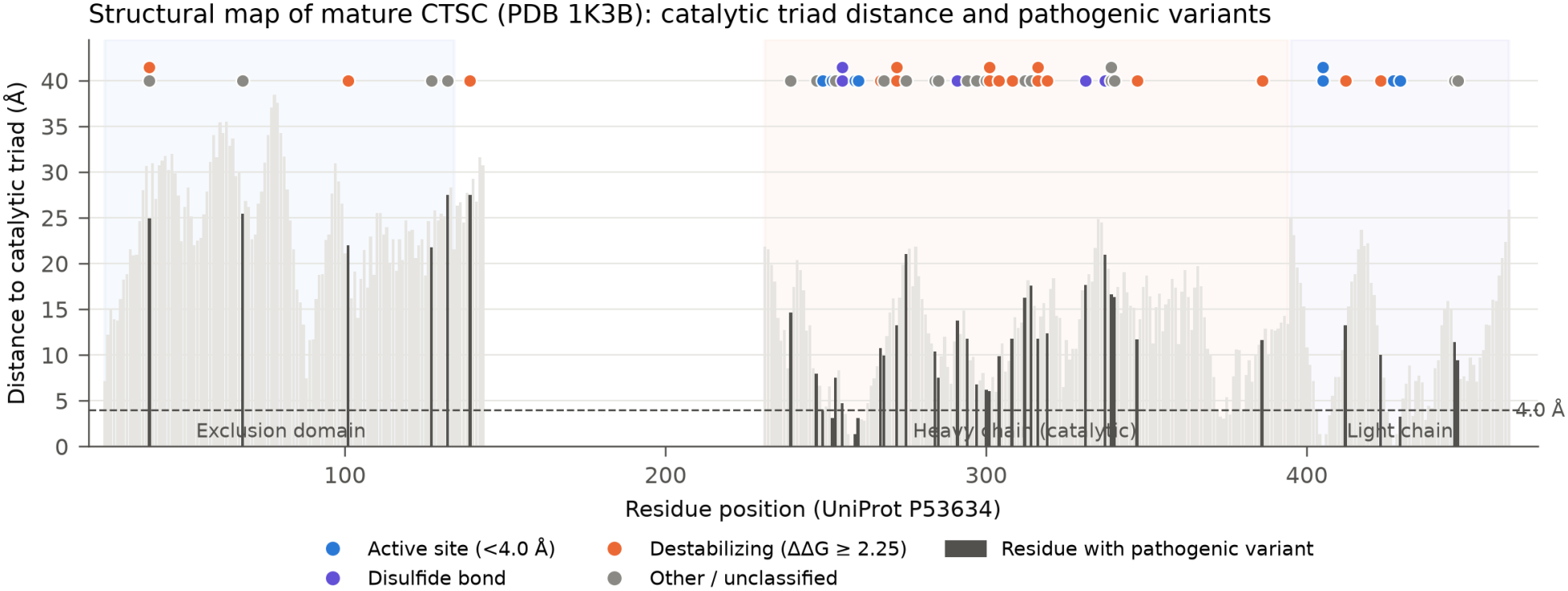
Structural map of mature human cathepsin C (PDB 1K3B). Minimum heavy-atom distance from each of the 352 resolved residues (UniProt P53634 numbering: exclusion domain 25–143, heavy chain 231–394, light chain 395–463) to the catalytic triad (Cys258, His405, Asn427). Dark gray bars indicate positions harboring pathogenic or literature-curated missense variants (n = 51 in 1K3B). Markers above bars classify each variant by primary structural mechanism: active site proximity (< 4.0 Å, blue), disulfide bond disruption (violet), thermodynamic destabilization (ΔΔG ≥ 2.25 kcal/mol, orange), or unclassified (gray).

**Figure 2.**
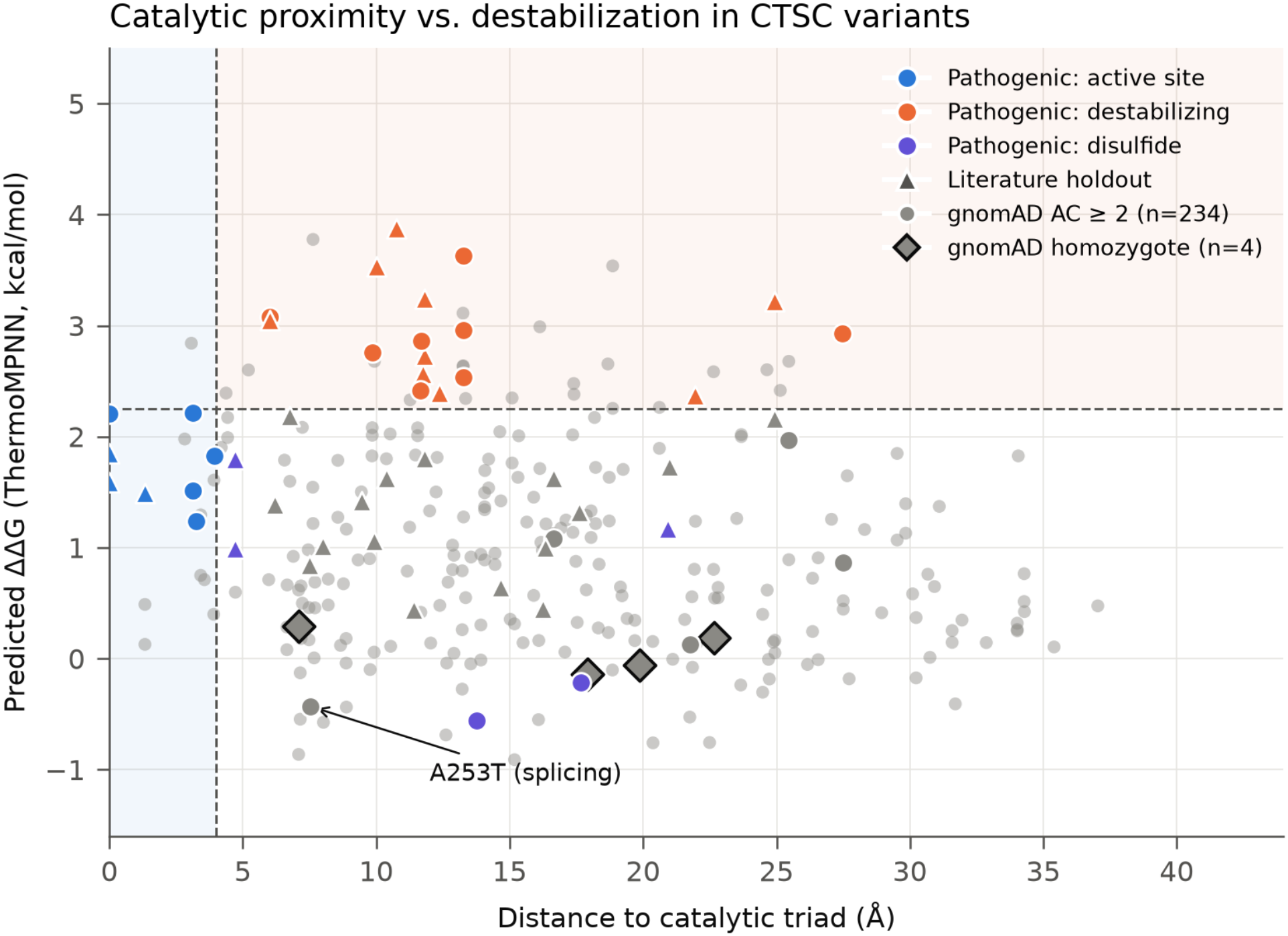
Catalytic triad proximity versus predicted thermodynamic destabilization (ΔΔG) across CTSC missense variants. Pathogenic variants (n = 51: calibration n = 20, circles; literature holdout n = 31, triangles) are plotted against gnomAD v4.1 recurrent population controls (AC ≥ 2, n = 234, gray circles) and homozygous controls (homozygote_count ≥ 1, n = 4, black-outlined diamonds). Dashed lines mark the pre-registered catalytic distance threshold (4.0 Å) and the leave-one-out calibrated stability threshold (2.25 kcal/mol). The exonic splicing variant A253T (c.757G>A, ΔΔG = -0.43 kcal/mol, 7.52 Å from catalytic triad) is highlighted as a protein-structure outlier rescued by AlphaGenome v2 splicing prediction (raw_score = 0.8589).

**Figure 3.**
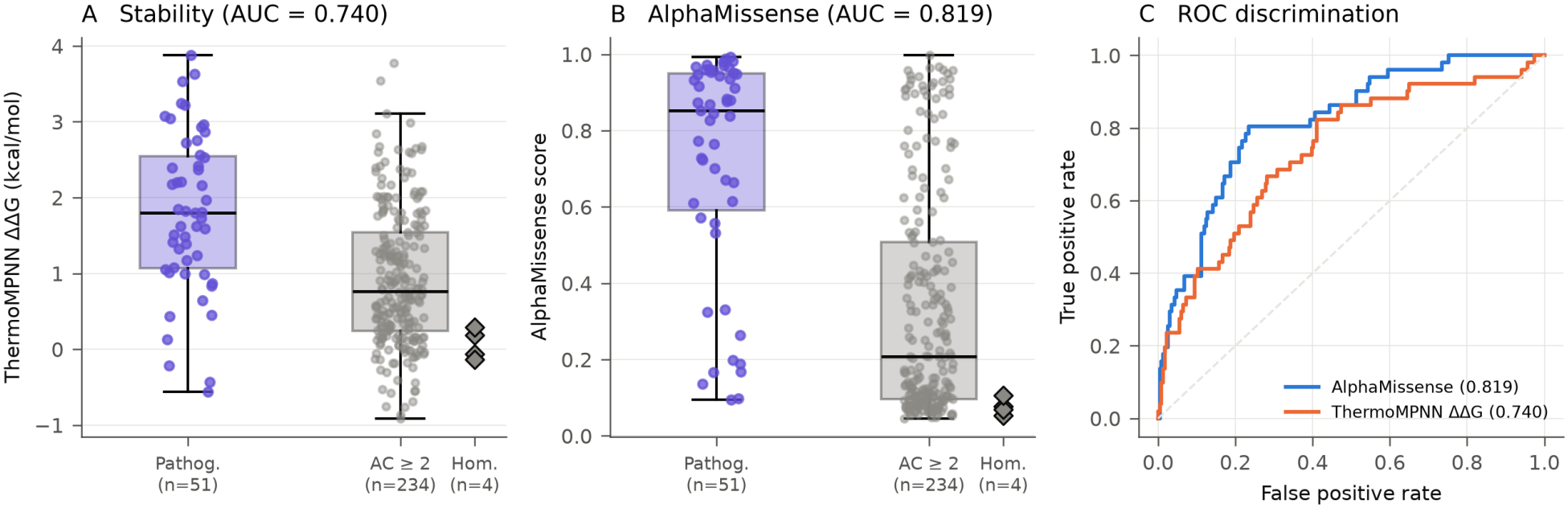
Discrimination between pathogenic CTSC missense variants and gnomAD v4.1 population controls. (A) ThermoMPNN ΔΔG (1K3B mature monomer): median 1.796 vs. 0.762 kcal/mol (AUC = 0.740, 95% CI 0.657–0.813, Mann-Whitney p = 8.33 × 10^−8^). (B) AlphaMissense pathogenicity score: median 0.853 vs. 0.208 (AUC = 0.819, 95% CI 0.753–0.877, p = 8.94 × 10^−13^). (C) Receiver operating characteristic (ROC) curves comparing AlphaMissense and ThermoMPNN ΔΔG. Homozygous gnomAD variants (n = 4) cluster near zero destabilization (-0.14 to 0.29 kcal/mol) and low AlphaMissense scores (0.053–0.106).

**Figure 4.**
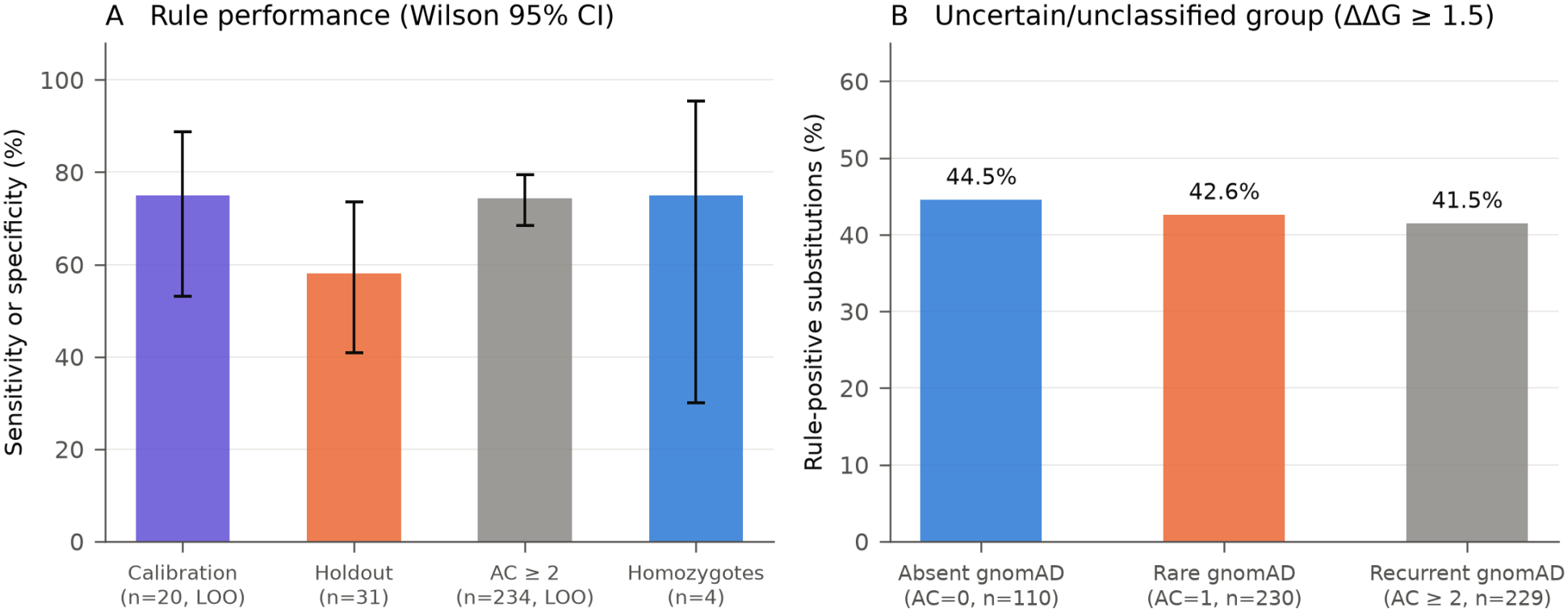
Rule validation and disjoint stratification of uncertain or unclassified substitutions. (A) Performance of the pre-registered four-term structural rule (ΔΔG ≥ 2.25 kcal/mol) with Wilson 95% confidence intervals across leave-one-variant-out calibration (15/20 = 75.0%), literature holdout (18/31 = 58.1%), AC ≥ 2 controls (174/234 = 74.4% specificity), and homozygous controls (3/4 = 75.0% specificity; 4/4 = 100% for the two-term rule). (B) Proportion of rule-positive substitutions in the residual group (ΔΔG ≥ 1.5 kcal/mol) stratified into mutually disjoint gnomAD subgroups: absent (AC = 0, 49/110 = 44.5%), singleton (AC = 1, 98/230 = 42.6%), and recurrent (AC ≥ 2, 95/229 = 41.5%). The recurrent residual group overlaps completely with the population comparators.

Applying the fixed modal threshold to the entire calibration positive set gave 17/20, an in-sample summary. In exploratory comparisons using the same folds, the two-term and stability-only rules gave 11/20 and 10/20 calibration positives, and 204/234 and 185/234 comparator negatives, respectively. They were positive for 12/31 and 14/31 holdout substitutions at their respective modal thresholds of 2.25 and 1.75 kcal/mol.

At 2.25 kcal/mol, the rule was positive for 18/31 substitutions in the historical literature holdout. The source audit found dated references through 2015 for 16 of these substitutions, of which 10 were rule-positive. Ten substitutions had only a reference traced after 2015, with 6 rule-positive at the fixed threshold. Five had unresolved dates, with 2 rule-positive. These counts are a description of the original assignments. Source chronology and case-level label verification would have to be established before evaluating temporal validation with the literature subset.

Using the four-term rule at 1.5 kcal/mol, positive fractions in the residual uncertain/unclassified group were 49/110 (44.5%) for substitutions absent from gnomAD, 98/230 (42.6%) for singletons, and 95/229 (41.5%) for recurrent substitutions. All 229 recurrent substitutions in this residual group also served as population comparators. These counts describe scores within the strata.

### 3.3 Functional observations and noncoding results

Four literature examples with reported DPPI activity deficiency were positive under the protein rule: p.Gln252Leu, p.Arg272Pro, p.Gly301Ser and p.Tyr347Cys (Pham et al., 2004). All four occurred in the calibration positives. This is descriptive agreement with selected reports. The p.Ala253Thr substitution, also in calibration, was negative under the protein rule (1K3B ΔΔG −0.43 kcal/mol; 7.52 Å to the catalytic triad). Published functional work on its c.757G>A allele found exon skipping and premature truncation (Abdel-Hamid et al., 2024). The AlphaGenome splice-site score of 0.8589 reported for this example is consistent with that RNA-level effect.

Of the 152 noncoding records, 12 carried pathogenic/likely pathogenic labels, 105 carried benign/likely benign labels and 35 formed the remaining group. All 12 positives were annotated as splice donor or acceptor variants, including one boundary-spanning deletion. None of the 105 negative-label records had those consequences. The consequence-only baseline and AlphaGenome splice-site score therefore each obtained AUC 1.000 in this comparison. Reported RNA-seq AUCs were 0.953, 0.955 and 0.967 for K562, HL-60 and fibroblasts, respectively, on the same allele set. The data lack a negative-label comparison group within the donor/acceptor annotations. Scores for deeper intronic candidates, including c.318+6826G>T, constitute exploratory predictions.

## 4. Discussion

The strongest pattern was the difference in predicted destabilization between positive and population-labelled substitutions. Catalytic proximity marked a smaller subset, and most structural features were weak discriminators at their tested cutoffs. A negative tetramer-interface result is consistent with limits of a proximity proxy; biochemical disruption can depend on the specific residue change, processing state or assembly context. The p.Ala253Thr case illustrates that a variant annotated as missense can affect transcript processing. The exploratory propeptide comparison suggests a question for follow-up in structures of procathepsin C.

Several features limit the interpretation. The original temporal grouping is unreliable. Several source references in the review table require allele-level adjudication. The positive-only holdout estimates sensitivity against a literature-derived label set and lacks an independent specificity estimate. Phenotype, segregation and functional evidence require variant-level review, particularly for substitutions supported only by literature. The recurrent comparator group may contain recessive carriers and includes unclassified substitutions. Cross-validation, bootstrap estimates, predictor coverage and the four homozygote comparators should be interpreted in light of those label definitions. The AlphaGenome contrast is fully separated by annotated consequence and the multi-lineage results use repeated scores on the same alleles. A consequence-matched benchmark and a comparison with an established splice predictor would address the incremental value of the model.

## Supporting information

Supplementary data, variant curation, and analysis code

## Data and code availability

The analysis scripts, consolidated inputs and outputs, and initial versioned analysis plan are available at https://github.com/contatofilipevono-rgb/papillon-lefevre-ctsc (audit baseline: f89ebaf797b188ffdc2621e2aa8bed13c5197b1b).

The supplementary materials include the 35-entry evidence ledger, the script reproducing the ClinVar-only comparison, and the historical exploratory figures.

## Supplementary figures · Exploratory analyses

**Figure 5.**
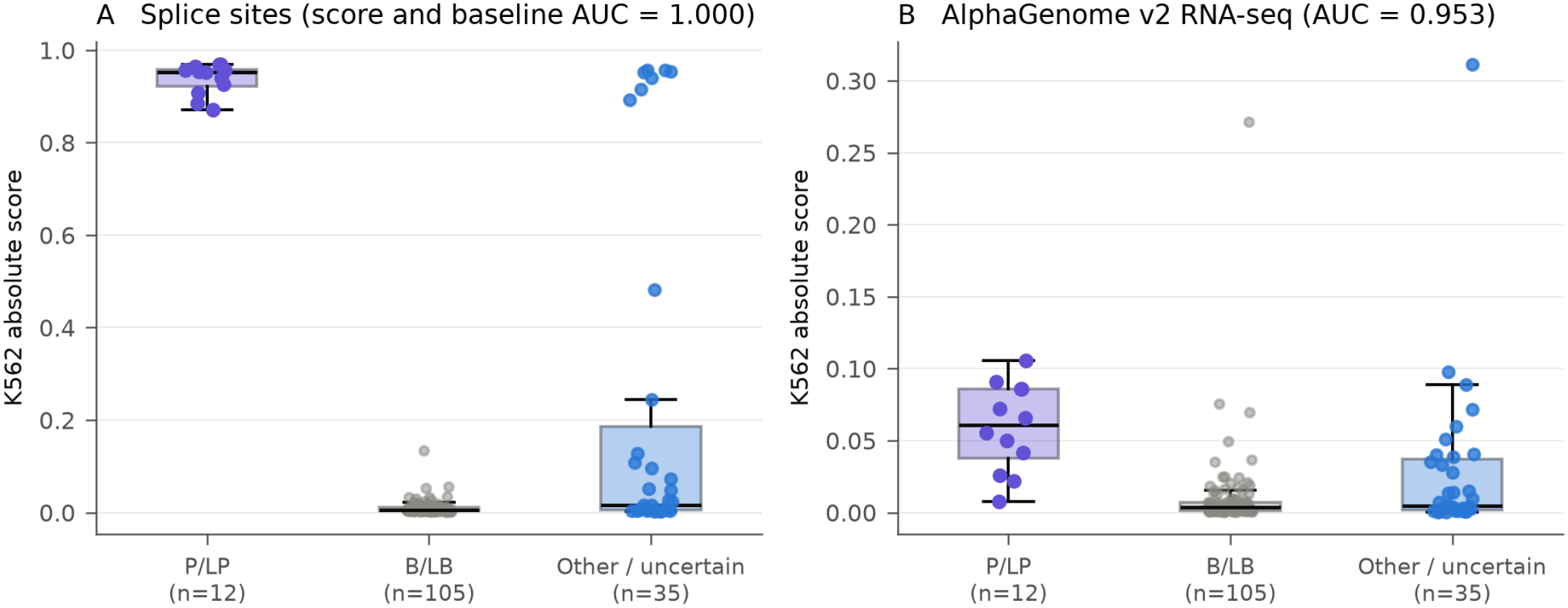
AlphaGenome v2 predictions for non-coding ClinVar CTSC variants (n = 152) in K562 myeloid cells (EFO:0002067). (A) Splicing (SPLICE_SITES) discriminates pathogenic/likely pathogenic (n = 12, median 0.953) from benign/likely benign (n = 105, median 0.006) variants (AUC = 1.000). An annotation-only splice donor/acceptor indicator produced the same AUC, because all 12 P/LP and none of the 105 B/LB records had those annotations. (B) Predicted RNA_SEQ impact (AUC = 0.953, p = 2.96 × 10^−7^). VUS (n = 35) include canonical splice variants and exploratory deep intronic candidates (c.318+6826G>T, 0.8932).

